# Comparative Analysis of Housing Temperature Impact on Heart Failure with Preserved Ejection Fraction in J vs N Strain C57BL/6 Mice

**DOI:** 10.1101/2024.08.05.606691

**Authors:** Rajesh Chaudhary, Tahra K. Suhan, Chao Wu, Afnan Alzamrooni, Ahmed Abdel-Latif

## Abstract

**Introduction:** Heart failure studies are conducted in preclinical animal models with different genotypic strains, 7-times higher metabolic rate, 5 to 6 times higher heart rate, and are housed in a cold-stressed environment of 23°C, unlike humans. These differences severely affect how animals respond to interventions, particularly those that lead to the development of metabolic syndrome, such as the two-hit model of diet-induced obesity (DIO) and N^μ^-nitro-L-arginine methyl ester (L-NAME) administration. A two-hit model of diet-induced obesity (DIO) and L-NAME administration has been proposed to induce heart failure with preserved ejection fraction (HFpEF) and mimic the hallmarks of metabolic syndrome and inflammation-induced heart failure in humans [1]. However, studies have reported conflicting results using this model. In this study, we examined the influence of mouse strain and environmental temperature on the development of metabolic syndrome and HFpEF using a two-hit model.

**Methods:** Eight-week-old, C57BL/6 mice (n=30) from the J and N strains were randomized to receive a high-fat diet (HFD) plus L-NAME versus a regular chow diet; and were randomized to be housed at a regular temperature of 23 °C versus a thermoneutral temperature of 30 °C. Glucose tolerance test (GTT, 2g/kg body weight), blood pressure via tail cuff, and echocardiography were conducted at baseline and, then at 5 and 15 weeks. Metabolic phenotyping was conducted at week 15 by using the Promethion Sable System.

**Results:** Our study revealed the significant effects of housing temperature and strain on the development of metabolic syndrome and HFpEF following the initiation of HFD +L-NAME over 5 and 15 weeks. At 5 weeks, both strains showed thermoneutral housing-induced attenuation of the effects of HFD + L-NAME on blood pressure and glucose tolerance, with the J strain exhibiting reduced diastolic dysfunction. By week 15, thermoneutral housing decreased energy expenditure (EE) and fat oxidation in both strains, while specifically reducing the respiratory exchange ratio (RER)_and glucose oxidation in J strain. Ejection fraction increased in both strains compared with the Chow group, except for J strain at 23 °C. Notably, physical activity levels remained constant across the groups, suggesting that the observed metabolic changes were not activity related. These findings highlight the complex physiological adaptations of these strains to different housing temperatures.

**Conclusions:** Thermoneutral housing conditions elicited strain-specific metabolic and cardiac effects in mice, with the J strain showing more pronounced responses. These findings highlight the critical influence of ambient temperature on experimental outcomes in rodent models, emphasizing the need to consider housing conditions when interpreting the results of metabolic and cardiovascular research.

## Introduction

Heart failure with preserved ejection fraction (HFpEF) is a complex clinical syndrome that accounts for nearly half of all clinical heart failure cases [2]. HFpEF is generally precipitated by comorbidities such as obesity, diabetes, and hypertension [3–7], resulting in systemic inflammation, a hallmark pathophysiology of HFpEF [8–10]. However, anti-inflammatory interventions have so far been unsuccessful [1]. Unsuccessful clinical interventions for HFpEF reflects our poor understanding of its pathophysiology and the lack of appropriate preclinical models.

Heart failure studies are conducted in preclinical animal models with different genotypic strains, approximately 7 times higher metabolic rate, and 5-6 times higher heart rate, and are housed in a cold-stressed environment of 23°C, unlike humans. These variations significantly impact how animals respond to metabolic stressors, leading to study results that do not accurately reflect human HFpEF conditions. This issue is further complicated by the diverse clinical presentations of HFpEF in humans [11]. Unlike in animal models, human HFpEF develops over time within a thermoneutral zone, defined as a temperature range of approximately 19-30 °C. This range represents the temperature at which a healthy individual can maintain a normal body temperature without increasing the energy expenditure (EE) above the basal metabolic rate [12]. However, the existing literature on HFpEF in animal models lacks consensus on the use of J [13] versus N strain C57BL/6 mice [14], including their optimal housing temperature. This study aims to develop a preclinical animal model of HFpEF that can faithfully replicate human HFpEF conditions to better understand and develop clinically translatable interventions. We deployed two-hit model, a coexistence of obesity and metabolic syndrome induced by HFD and mechanical stress induced by L-NAME-mediated nitric oxide (NO) synthase suppression to develop HFpEF phenotype [14].Using the two-hit model, this study compares the effects of J vs. N strain mice and housing temperatures (standard 23 °C vs. thermoneutral 30 °C) on the development of metabolic syndrome and HFpEF.

C57BL/6J mouse has traditionally been preferred as a preclinical DIO-induced disease models due to a mutation in nicotinamide nucleotide transhydrogenase (*Nnt*) gene that predisposes it metabolic syndrome phenotype [15]. Spontaneous mutation of nicotinamide nucleotide transhydrogenase located in the inner mitochondrial membrane contributes to a decrease in the NADPH/NADP(+) ratio, resulting in mitochondrial redox imbalance [16] that drives metabolic syndrome, systemic inflammation [17] and heart disease [18]. Additionally, cold-stressed mice at a standard housing temperature of 23 °C tend to increase their total energy expenditure (EE) by 30% through non-shivering adaptive thermogenesis to maintain core body temperature [19]. Therefore, we hypothesize that the increased predisposition of the J strain to diet-induced obesity and metabolic phenotype makes it an ideal candidate for the DIO HFpEF mouse model over the N strain as they are genetically better equipped to compensate for energy lost due to adaptive thermogenesis under standard housing conditions without compromising the DIO phenotype required for HFpEF. In this study, although thermoneutral housing reduces the metabolic phenotype in both strains equally, the indices of heart failure were more impacted in the J strain at thermoneutral housing due to greater fluctuation in HFpEF phenotype. This study, therefore, stabilizes the J strain at 23 °C as a better predictive model for HFpEF over the N strain, unlike previously observed [14].

## Research design and methods

### Animals

All animal experiments were performed according to the approved University of Michigan Institutional Animal Care and Use Committee and the Institutional Animal Care and Use Committee (IACUC) protocol. Eight-week-old C57BL/6 mice of both the J and N strains (n=30, Jackson Laboratory, Bar Harbor, ME) were included in this study. Mice underwent a two-week acclimatization period at an ambient temperature of 23 °C prior to randomization to one of the six groups that included two different housing temperature conditions (23 °C and 30 °C): (1) Strain J, receiving standard Chow at 23^0^C (n=3); (2) Strain N, standard Chow at 23^0^C (n=3); (3) Strain J, high-fat diet (HFD, #D12492, Research Diet, USA) supplemented with nitric oxide synthase inhibitor, *N*^ω^-nitro-L-arginine methyl ester (L-NAME, 0.5 g/L or 100 mg/200 ml) (#328750050, Thermo Scientific, USA) in drinking water at 23 ^0^C (n=6); (4) Strain N, HFD with L-NAME at 23 °C; (5) Strain J, HFD with L-NAME at 30 °C; and (6) Strain N, HFD with L-NAME at 30 °C. All mice were housed 2-3 mice/cage in 12 h of light:12 h of darkness with lights off at 07:00 (Zeitgeber, ZT 0) and lights on at ZT 12 and had ad libitum access to food (Chow, 18% calorie from fat and HFD, 60% calorie from fat) and water (normal for Chow group and L-NAME containing for HFD+L-NAME group) throughout the 15-week study duration. Food, water, and cages were changed twice weekly for all mice including the Chow group to standardize the handling at ZT14. Heart, liver, spleen, whole blood, total bone marrow, urine, and stool were snap-frozen in liquid nitrogen and stored at -80 °C for further processing. Approximately 50-70 mg of ventricle was separated from the whole heart on an ice-cold metal plate before freezing briefly (2-3 sec) in semisolid Isopentane placed on dry ice and immediately snap-frozen in liquid nitrogen in cryovials, and later transferred to -80 °C for further metabolomics study. Whole blood was collected via direct cardiac puncture, placed in tubes containing CTAD at 1:10 and plasma samples were prepared after spinning at 700g for 5 minutes at room temperature. The supernatant was collected and centrifuged again at 10,000g for 10 minutes at room temperature. Bone marrow was collected from the tibia and femur of the two hind legs of mice as previously described [20].

### Intraperitoneal glucose tolerance test (IPGTT)

Intraperitoneal glucose tolerance test (IPGTT) was performed at weeks 5 and 15 after 5h of fasting using 2 g/kg body weight (3-6 mice/group). Blood glucose readings were measured at 0, 15, 30, 60, 90 and 120 minutes following the bolus dose via tail-vein prick using Accu-Chek® Performa II, Roche, USA). The area under the curve (AUC) was calculated using the trapezoidal rule [21].

### Body weight and composition

Body weight was recorded weekly during cage, food, and water change (ZT14). Body composition was measured in a subset of mice (n=3/group) using an NMR-based analyzer EchoMRI^TM^-500 Body Composition Analyzer, according to manufacturer’s instructions, at the University of Michigan Phenotyping Core.

### Sable Metabolic cage studies

At week 13, a subset of mice (n=3/group) was acclimatized for a week to single housing conditions at their respective 23 °C and 30 °C housing temperatures before transferring to Promethion (Comprehensive, High-Resolution Behavioral Analysis Systems, Sable Systems International, USA) maintained at 23^0^C and 30^0^C. Food and water consumption and x, y, and z beam breaks, VO2 and VCO2, were measured at 20-minute intervals. Energy expenditure (EE) and respiratory quotient (RQ) were calculated using the Weir equation [22]. EE was adjusted to raw EE/ (body weight)^3/4^ as previously described [23]. The University of Michigan Animal Phenotyping Core also performed these tasks.

### Blood pressure measurements

Blood pressure measurements were conducted using a CODA Monitor (Kent Scientific) via the tail-cuff method, according to the manufacturer’s instructions, after the mice were acclimatized and settled in a quiet procedure room with no disturbance. Briefly, an occlusion (O) tail cuff and a volume pressure recording (VPR) cuff were placed on the tail of the mouse, and the O-cuff was inflated to impede blood flow to the tail before the cuff was deflated slowly and return blood flow was measured by using a VPR sensor. Measurements were collected after ensuring the body temperature was between 32 °C and ≤37 °C. Body temperature was recorded at the base of the tail using a laser temperature gun. All measurements were recorded in non-anesthetized mice and body temperature was maintained using a thermal pad.

### Echocardiography measurements

All echocardiographic measurements were recorded in anesthetized mice maintained with 1-2% isoflurane and 95% oxygen to maintain the heart rate at 450±50 beats per minute. Briefly, trans- thoracic echocardiography was conducted using the Vevo F2 Imaging System (#53699-20, FUJIFILM VisualSonics, Inc., USA). The anterior chest hair was removed using Nair Hair-removal cream before sedation. Body temperature-maintained pre-warmed ultrasound gel was applied to the area underlying the heart after immobilizing the mice on the stage. The parasternal short-axis view, indicated by the presence of papillary muscles was used to obtain the M mode for ejection fraction, fractional shortening, left ventricular mass, cardiac output, and other cardiac parameters. The apical four-chamber view was used to obtain the tissue Doppler and mitral valve pulse-wave Doppler measurements for myocardial tissue and blood flow velocity, respectively. All parameters were measured at least three times or more times, and all individual measurements were plotted using their means for statistical analysis.

## Statistical analysis

Data are presented as mean± SEM. The data were checked for normality using the Shapiro-Wilk test, and all statistical analyses were performed on log-transformed data if not normally distributed using either SPSS (version 29.0.2.0, IBM, Armonk, NY, USA) or GraphPad Prism (Version 10.2.2 (341)). One-way ANOVA was performed for group comparison after separating the data based on mouse strains. *P*<0.05 was considered statistically significant.

## Results

### Thermoneutral housing temperature reduces the indices of metabolic phenotype in the DIO-HFpEF mouse model in C57BL/6J and C57BL/6N strains

Considering the challenges associated with recapitulating human HFpEF in animal models, we utilized the two-hit model of DIO-induced HFpEF mouse model with J and N strains housed at 23 °C and 30 °C to examine the effect of housing temperature and strain background on the development of HFpEF. Initially, our study aimed to ascertain the impact of two distinct housing conditions on the metabolic phenotype of mice and the development of HFpEF. This interest was prompted by observations that mice housed at 23 °C exhibit a 30% increase in energy expenditure (EE) to maintain core body temperature via lipolysis in adipocytes [24], potentially influencing the metabolic ramifications in the two-hit HFpEF model [19]. High-fat diet and L-NAME-induced HFpEF, hereafter referred as (DIO-HFpEF) was induced by following the two-hit model [14]. Although HFpEF is clinically presented with great heterogeneity [11], it is commonly precipitated by obesity, elevated arterial hypertension, and reduced glucose tolerance, a phenotype widely referred to as metabolic syndrome [25, 26]. As expected, the J strain DIO mice housed at 23 °C and 30 °C gained significant weight at week 15 compared to the Chow group **(Figure 1A)**, whereas the N strain DIO mice at 30 °C gained significantly more weight compared to both Chow and DIO mice at 23 °C **(Figure 1E)**. The J and N strain DIO mice at 30 °C consumed considerably fewer calories compared to Chow-fed mice at 15 weeks **(Figure 1B** and **1F, respectively)**, while only the J strain DIO at 30 °C consumed fewer calories compared to DIO at 23 °C **(Figure 1B)** as did was the N strain DIO at 23 °C compared to Chow **(Figure 1F)**. However, no difference was observed between the 23 °C vs 30 °C housed N strain DIO mice. Percentage of body fat was higher in both J and N strains housed at 23 °C **(Figure 1C** and **1G**, respectively) and 30 °C **(Figure 1C** and **1G, respectively)** than in the Chow diet, with no significant interaction with strains or housing temperature. The percentage of lean mass in both the DIO J and N strains at 23 °C and 30 °C was significantly lower than in the Chow diet **(Figure 1D** and **1H)**. Blood pressure measurements at Week 5 revealed significantly elevated systolic blood pressure in J strain only **(Figure 1I)** while both systolic and diastolic blood pressures were elevated in N strain at 23 °C **(Figure 1K** and **1L)**. The diet-induced effect on the systolic blood pressure was attenuated in N strain housed at 30 °C compared to those housed at 23 °C but was still significantly elevated compared to Chow-fed animals **(Figure 1K)**. Diastolic blood pressure in the N strain DIO was reduced in 30 °C housed mice compared with that in mice housed at 23 °C **(Figure 1L)**, while no difference was observed between DIO at 30 °C vs Chow-fed animals **(Figure 1L)**. At week 5, both J and N strain DIO mice housed at 30 °C had less pronounced glucose intolerance compared to their DIO counterparts at 23 °C **(Figure 1Q** and **1S)**, and Chow-fed animals **(Figure 1Q** and **1S)**. At week 15, glucose tolerance was decreased in both the J and N strain DIO mice housed at both housing temperatures compared to Chow-fed animals. However, this effect was more pronounced in the J strain **(Figure 1R** and **1T)**. These results suggest important effects of inbred mouse strain and housing temperature on the development of metabolic syndrome in the DIO-HFpEF model.

**Figure 1.**
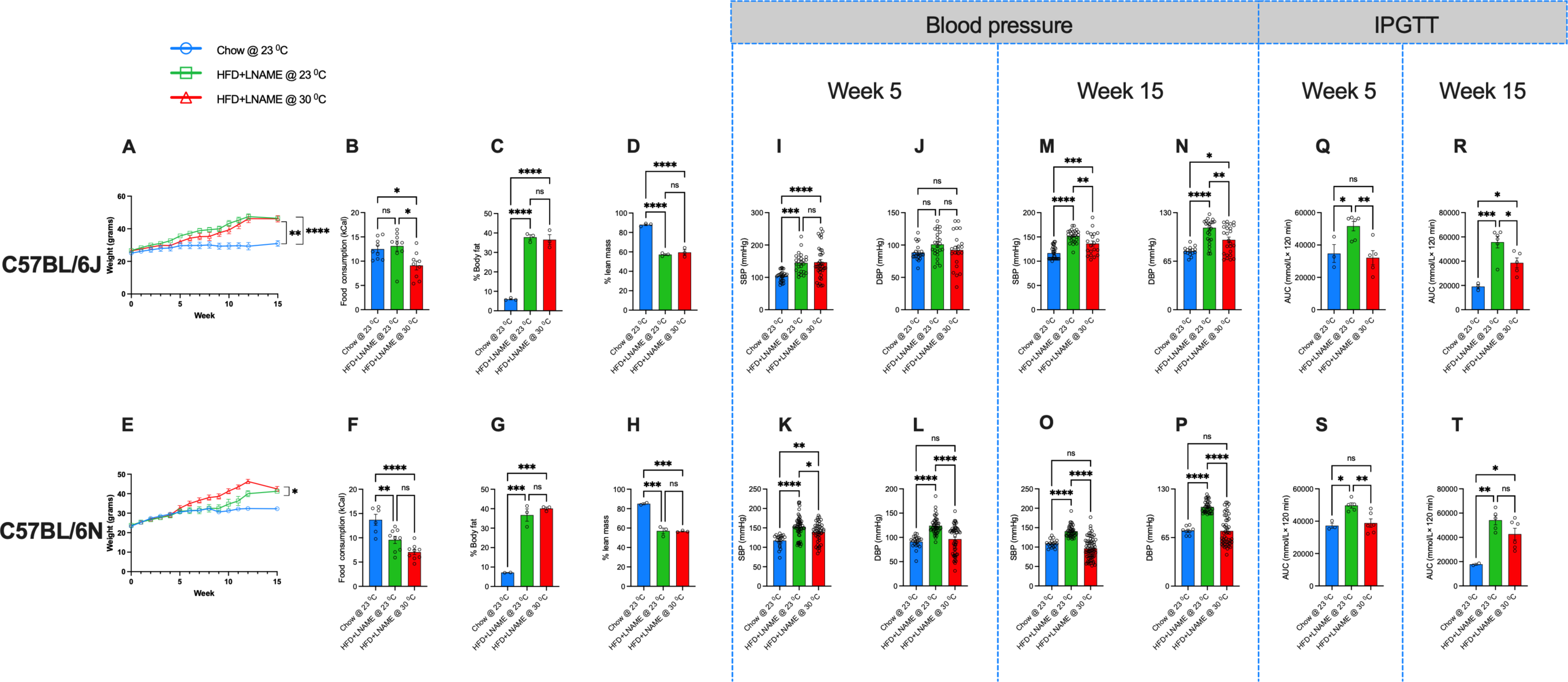
(A-D) Weekly weight gain, food consumption (kCal), percentage body weight, and percentage lean mass for J strain, and **(E-H)** for N strain, respectively. **(I, J** and **M, N)** Systolic and diastolic blood pressure measurements at week 5 and week 15, respectively, for J strain. **(K, L** and **O, P)** Systolic and diastolic blood pressure measurements, for N strain. **(Q, R)** Week 5 and week 15, intraperitoneal glucose tolerance test (IPGTT, 2g/kg) for J strain and **(S, T)** for N strain, respectively. HFD: High-fat diet, L-NAME: N^μ^-nitro-L-arginine methyl ester, SBP: systolic blood pressure, DBP: diastolic blood pressure, %: percentage, kCal: kilo calorie, AUC: Area under the curve, IPGTT: intraperitoneal glucose tolerance test. One-way ANOVA with Tukey’s *post hoc* test. *P<0.05, **P<0.01, ***P<0.001, and ****P<0.0001, considered as statistically significant. N=3-6 mice/group.

### Housing temperature altered echocardiographic indices of diastolic dysfunction in both C57BL/6 mouse strains at 5 weeks

Thermoneutral housing condition of 30 °C is reported to worsen systolic function in transverse aortic constriction in mice [24]. Our study is the first to report the effects of thermoneutral housing conditions on the development of DIO-HFpEF. Interestingly, 5 weeks of HFD with L-NAME induced the DIO-induced HFpEF phenotype only in the J strain mice but not in the N strain mice, regardless of the housing temperature conditions (23 °C or 30 °C), contrary to a previous observations [14]. Only the J strain DIO HFpEF housed at 30 °C had diastolic dysfunction as evidenced by the increased E/A and E/E’ ratios compared to those of the 23 °C DIO and Chow-fed groups **(Figure 2B and C**). As expected, ejection fraction (EF) was not different between the groups in either strain **(Figure 2A** and **G)** at week 5. The fraction shortening percentage remained unchanged in the J strain between the groups but was marginally decreased in the N strain DIO at 23 °C compared to Chow **(Figure 2D** and **J)**. Similarly, we did not observe a significant effect of DIO, housing temperature, or mouse strain on the left ventricular end-systolic and end-diastolic diameters (**Figure 2K** and **E**, respectively).

**Figure 2.**
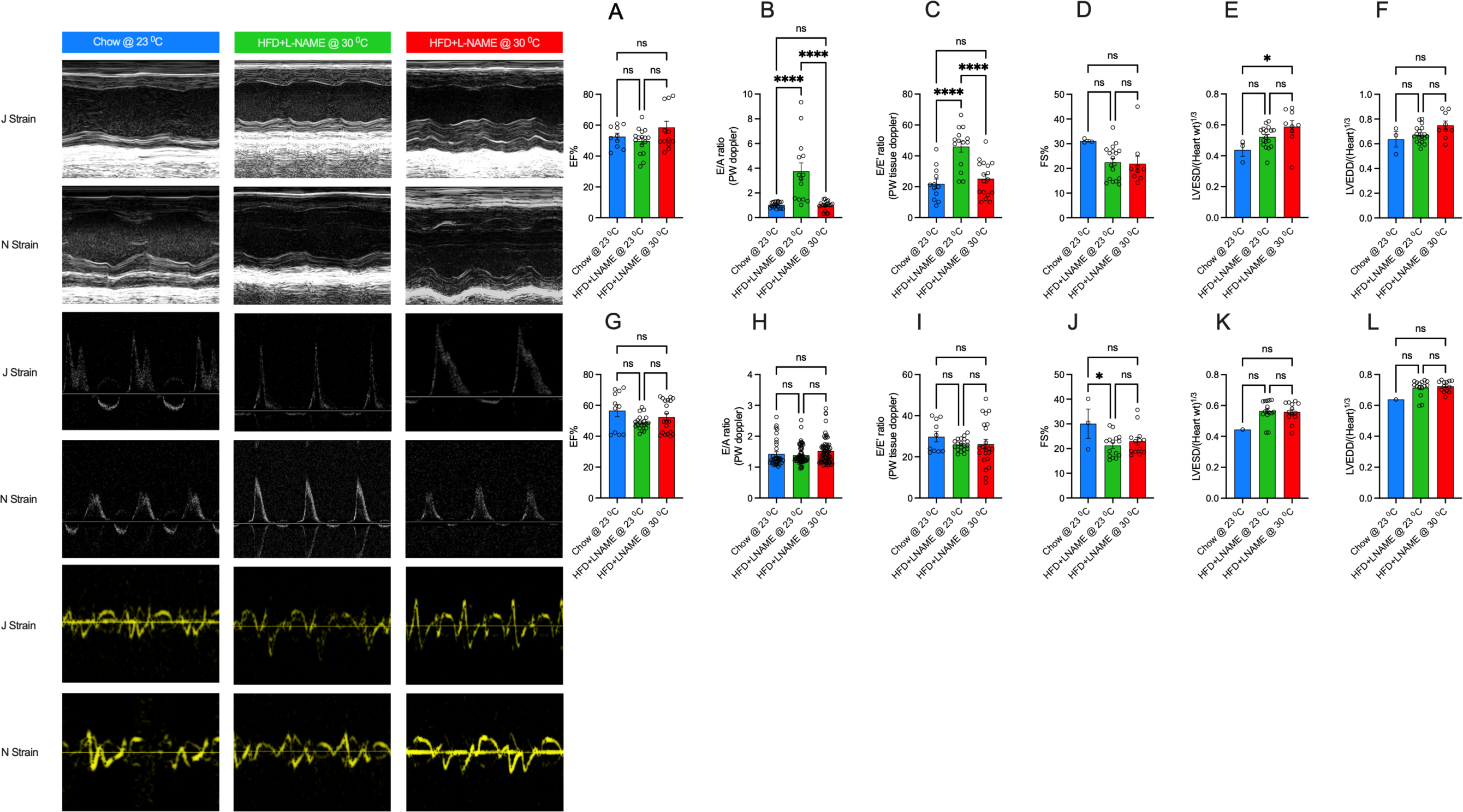
(A-F) Percentage ejection fraction, pulse-wave doppler, tissue doppler, percentage fractional shortening, left ventricle end systolic diameter, left ventricle end diastolic diameter for J strain, and **(G-L)** for N strain **at week 5,** respectively. EF%: percentage ejection fraction, FS%: percentage fractional shortening, LVESD: left ventricle end diastolic diameter, LVEDD: left ventricle end diastolic diameter, PW: pulse wave. One-way ANOVA with Tukey’s *post hoc* test. *P<0.05, **P<0.01, ***P<0.001, and ****P<0.0001, considered as statistically significant. N=3-6 mice/group.

### Thermoneutral housing differentially impacts indices of diastolic function at week 15 in both C57BL/6 strains

At Week 15, the E/A and E/E’ ratios increased regardless of the strain or housing temperature compared to Chow (Figure 3B and **H**). The ejection fraction percentage in both strains increased in the DIO model at 30 °C compared with Chow and their DIO counterparts at 23 °C (Figure 3A and **G**) but only the N strain DIO at 23 °C had an increased ejection fraction percentage compared with that of Chow, with no such difference observed in the J strain (Figure 3A and **G**). A subtle increase in the ejection fraction percentage for both strains at 30 °C including an increase in N strain DIO at 23 °C (Figure 3A and **G**), which is indicative of the aging effect as observed in clinically diagnosed HFpEF patients [27]. Fractional shortening did not changed across the groups in the J strain; however, it was increased in the N strain DIO at both housing conditions (Figure 3D and **J**). The left ventricle end-systolic diameter remained unchanged across the groups in both strains, but the end-diastolic diameter was slightly decreased in the J strain DIO at 23 °C compared to Chow **(Figure 3F)**.

**Figure 3.**
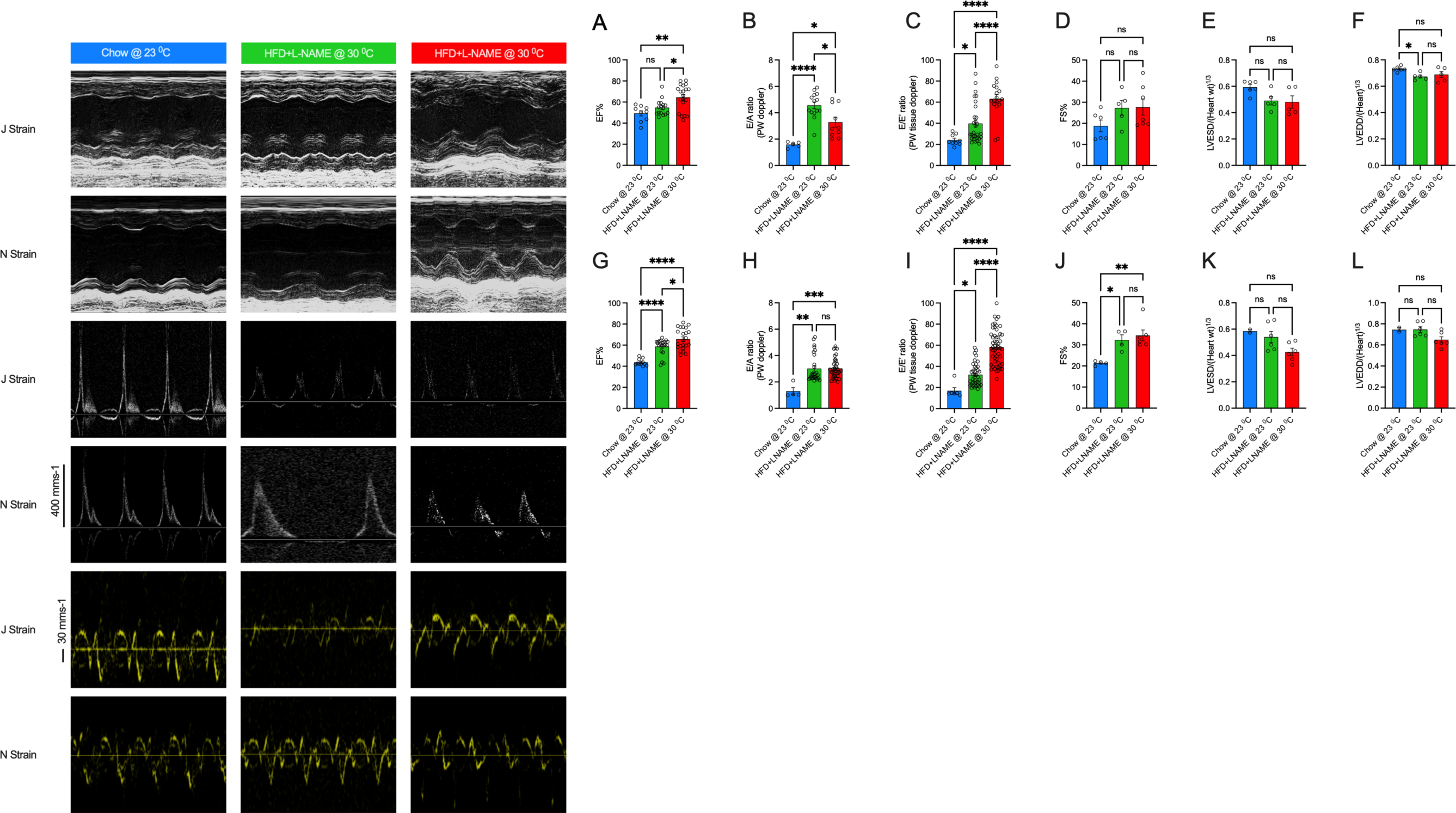
(A-F) Percentage ejection fraction, pulse-wave doppler, tissue doppler, percentage fractional shortening, left ventricle end systolic diameter, left ventricle end diastolic diameter for J strain, and **(G-L)** for N strain **at week 15,** respectively. EF%: percentage ejection fraction, FS%: percentage fractional shortening, LVESD: left ventricle end diastolic diameter, LVEDD: left ventricle end diastolic diameter, PW: pulse wave. One-way ANOVA with Tukey’s *post hoc* test. *P<0.05, **P<0.01, ***P<0.001, and ****P<0.0001, considered as statistically significant. N=3-6 mice/group.

### Thermoneutral housing temperature alters metabolic rate/activity and energy expenditure in both C57BL/6 mouse strains

Our studies showed that a 30 °C thermoneutral housing temperature attenuated the effect of DIO on the development of metabolic syndrome in a C57BL/6 mouse model. This was evidenced by the reduced effect of DIO on glucose tolerance, blood pressure, and body weight gain compared to animals housed at 23^0^C. Additionally, energy expenditure was lower in DIO HFpEF mice at 30 °C, suggesting that thermoneutral housing affects metabolic phenotypes associated with DIO-induced HFpEF. However, as previously reported [28], these mice also had reduced total calorie consumption when housed at 30 °C compared to animals housed at 23 °C and the Chow-fed group. Since these metabolic phenotypes play a critical role in the pathophysiology of DIO- HFpEF, we further conducted a SABLE metabolic chamber experiment to determine the underlying mechanisms between the metabolic and echocardiographic differences observed at the different housing temperatures.

Interestingly, the energy expenditure (EE) was lower in DIO-HFpEF mice at 30 °C than in both Chow and their DIO counterparts at 23 °C in both the active and inactive phases **(Figure 4A-F)**. Respiratory exchange ratio (RER) was decreased in both the J and N strains at both 23 °C and 30 °C in both the active and inactive phases compared to Chow (**Figure 4G-L)**. RER was further decreased in 30 °C housed mice but only in the J strain in both active and inactive phases **(Figure 4G-I)**. RER of 0.7 indicates fat metabolism as the sole source of energy, while a ratio of 1.0 indicates carbohydrate metabolism as the main source of energy. As expected, both strains of DIO at both 23 °C and 30 °C had higher fat oxidation than Chow in both active and inactive phases **(Figure 4S-X)**. Interestingly, however, fat oxidation was still lower in the 30 °C DIO of both strains, even though their RER was lower than their DIO counterparts at 23^0^C in both strains. This observation was independent of their physical activity, which was lower in both DIO-HFpEF at 23 °C and 30 °C **(Figure 4Y-Z4)**. These results suggest that thermoneutral temperature mitigates some of the adverse effects of DIO on metabolic health, such as improving glucose tolerance and reducing body weight gain. Reduced energy expenditure and altered metabolic phenotypes at this temperature may contribute to these protective effects. The RER findings suggest a shift towards increased fat oxidation under DIO conditions, especially notable in the thermoneutral housing scenario.

**Figure 4.**
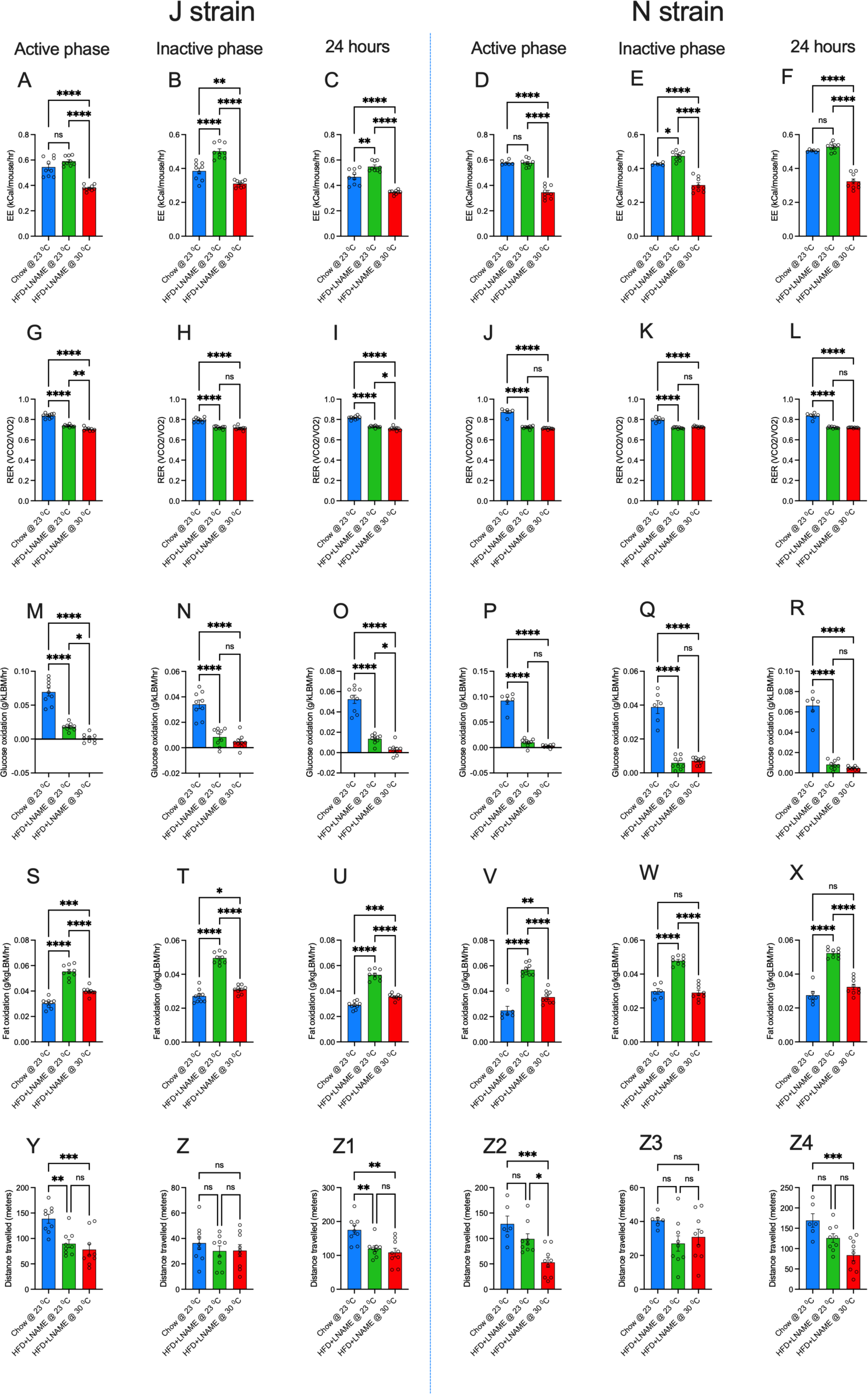
(A, G, M, S, and Y) Energy expenditure, respiratory exchange ratio, glucose oxidation, fat oxidation, and distance travelled in active phase, **(B, H, N, T,** and **Z)** in inactive phase, and **(C, I, O, U,** and **Z1)** in 24 hours, respectively, for J strain. **(D, J, P, V,** and **Z2)** Energy expenditure, respiratory exchange ratio, glucose oxidation, fat oxidation, and distance travelled in active phase, **(E, K, Q, W,** and **Z3)** in inactive phase, and **(F, L, R, X,** and **Z4)** in 24 hours, respectively, for N strain. EE: energy expenditure, RER: respiratory exchange ratio. VCO2: volume of carbon dioxide exhaled, V02: volume of oxygen inhaled. One-way ANOVA with Tukey’s *post hoc* test. *P<0.05, **P<0.01, ***P<0.001, and ****P<0.0001, considered as statistically significant. N=3-6 mice/group.

## Discussion

Heart failure with preserved ejection fraction (HFpEF) is marked by considerable clinical heterogeneity, posing significant challenges in accurately mimicking this condition in preclinical models subsequently hindering the development of effective clinical interventions [7, 29]. Commonly, preclinical mouse studies are conducted at standard housing temperatures ranging from 20-26^0^C, conditions under which mice exhibit profound physiological and metabolic changes such as increased total energy expenditure [30] and a nearly 2-fold increase in heart rate [31]. This study compares the effects of mouse strain and standard vs. thermoneutral housing conditions in DIO-induced HFpEF in the two-hit model. As previously observed [24], thermoneutral housing attenuated the impact of DIO on both systolic and diastolic blood pressure and improved glucose tolerance compared with standard housing temperature [24]. Thermoneutral temperature reduced the total energy expenditure only in the J strain, possibly by reducing the fat oxidation. These metabolic changes resulted in more consistent echocardiographic diastolic dysfunction findings in the J strain housed at standard room temperature. Our findings suggest that the mouse strain and housing temperature play important roles in the development of metabolic syndrome and diastolic dysfunction in the C57BL/6 mouse models. These factors significantly influenced the experimental findings in DIO-based HFpEF models, highlighting the importance of considering housing conditions and strain selection in future studies.

The current study was conducted to examine the role of the C57BL/6 strain and housing temperature on mouse response to diet-induced obesity. We attempted to explain some of the observed variability in the results of preclinical HFpEF studies. Preclinical mouse studies are conducted at a standard housing temperature of 20-26^0^C that subjects mice to a constant cold stress [32], unlike their habitat’s thermoneutral temperature of 28-32^0^C [33]. However, even a modest change in temperature profoundly impacts the systemic energy balance [30], including total calorie consumption [19], heartbeat [28], and physical activities [34], which influence the experimental findings in preclinical mouse models such as DIO-HFpEF [24]. Although previous studies have reported the effects of housing conditions on murine metabolic phenotypes, to our knowledge, this is the first study that reports the effects of thermoneutral housing on the development of diastolic dysfunction using the two-hit model.

Our findings suggest that thermoneutral housing attenuates the overall metabolic burden associated with DIO equally in both C57BL/6 strains independent of physical activity as evidenced by lower blood pressure and better glucose tolerance. Some of these findings are consistent with a previous observations [35]. Cold-stress is known to increase the circulating level of catecholamine [24, 36, 37] which drives the hypertensive response [38] and predisposes the patient to cardiomyopathy [39, 40]. Indeed, cold-induced catecholamine triggers the blood pressure increase by 5.5 to 6.5 mmHg with every 10 °C drop in temperature [41], supporting our findings that thermoneutral housing lowers blood pressure compared to standard housing temperature. Although the findings of thermoneutral housing condition may suggest improved metabolic burden during DIO, it is worth noting that thermoneutral housing may uncouple metabolic inflammation in adipose tissue from the development of insulin resistance [42], as well as DIO-induced cardiovascular complications such as atherosclerosis [42]. As observed at the metabolic front, thermoneutral housing also improved the indices of diastolic dysfunction as evidenced by the lower E/A and E/E’ ratios at week 5. However, these effects did not last until week 15, and both housing temperatures manifested with a comparable degree of diastolic dysfunction. This observation aligns with the impact of two-hit model’s on the HFpEF indices, which appears to override the initial benefits of thermoneutral housing. The progression of diastolic dysfunction over time, regardless of housing temperature, suggests that the DIO-induced cardiovascular complications may eventually develop despite initial improvements in metabolic parameters [19, 24]. Along the same trend and contrary to previous observations [19], we observed a reduction in the respiratory exchange ratio, which corroborated the reduction in both fat and glucose oxidation, independent of physical activity. Despite the decrease in fat oxidation under thermoneutral conditions, the lower respiratory exchange ratio indicates that the mice still relied more heavily on fat as their primary energy source, which explains their reduced weight gain under thermoneutral housing conditions. A lower respiratory exchange ratio is considered as an auxiliary physical fitness indicator in trained subjects [43, 44], and a single bout of aerobic and resistance exercise reduces RER for at least 24 hours post-exercise [45]. It is worth noting that reduced RER in thermoneutral housed mice may confer health benefits to the heart, which derives 95% of its energy from oxidative phosphorylation [46].

Our study has some limitations, including a small sample size and reliance on male mice. We opted to include only male mice in our studies based on observations demonstrating that female mice are resistant to this two-hit model of HFpEF [47]. Due to logistic limitations, measurements such as GTT, blood pressure and echocardiography were conducted at the standard housing temperature for all groups, including the thermoneutral-housed mice. Therefore, future studies should include sex and temperature-matched groups to eliminate confounding factors that may have influenced the findings of this study.

In conclusion, our findings suggest that the housing temperature and strain selection in C57BL/6 mouse models significantly influence the experimental results of DIO-based HFpEF studies. These factors contribute to variations in metabolic responses, blood pressure, and cardiac diastolic function indices, highlighting the importance of considering housing conditions in future preclinical research. Collectively, this study indicates that the C57BL/6J strain housed under standard housing conditions developed a more consistent HFpEF phenotype when challenged with the two-hit metabolic model.

## Acknowledgments

This study was supported by grants from the National Institutes of Health (R01 grant number HL124266) and the VA Merit award #I01- CX002684-01.

## Authors’ contributions

RC planned the project, performed experiments, analyzed data, and wrote the manuscript. TKS and AA helped manage the project, collected data, and revised and edited the manuscript. CW performed the echocardiography on the mice, helped collect the data, analyze and interpret results, and revised the manuscript. AAL conceptualized the project, wrote, revised, and edited the manuscript. All authors revised and edited the manuscript.

## Conflict of interest

Authors declared no conflict of interest.

## Notes

### Competing Interest Statement

The authors have declared no competing interest.

